# Temporal Microbial Community Dynamics within a Unique Acid Saline Lake

**DOI:** 10.1101/2020.12.17.423355

**Authors:** Noor-ul-Huda Ghori, Michael. J. Wise, Andrew. S. Whiteley

## Abstract

Lake Magic is one of the acidic hypersaline lakes (ca. 1 km in diameter) present within the Yilgarn Craton in WA. This unique lake exhibits extremely low pH (<1.6) coupled to very high salinity (32% TDS) with the highest concentration of aluminium (1774 mg/L) and silica (510 mg/L) in the world. Previous studies on Lake Magic diversity has revealed that the lake hosts acidophilic, acidotolerant, halophilic and halotolerant bacterial species. These studies provide indicators of the population residing within the lake. However, they do not emphasize the survival mechanisms adopted by the resident microorganisms and how the diversity of microbial populations residing within the lake changes during the dynamic stages of flooding, evapo-concentration and desiccation. We have studied the bacterial and fungal diversity in Lake Magic via amplicon sequencing and functional analysis through different stages of the lake in a span of one year, in the salt and sediment layer. Our results highlight that the diversity in Lake Magic is strongly driven by the pH and salt concentrations at different stages of the lake. The microbial community becomes more specialised in specific functions during more extreme stages. This also suggests that microbial interactions are involved in stabilising the ecosystem and is responsible for the resistance and resilience of these communities as the interactions of these microbes create a safe haven for other microbes to survive during more extreme stages.

## 1 Introduction

Acid saline lakes represent one of the most extreme aquatic environments on Earth. They are poly-extreme ecosystems, exhibiting extremely acidic pH and salinities close to saturation. Such environments are of significant microbial interest as they host organisms that are not only capable of withstanding pH and salinity stress, but also survive in the presence of additional stressors such as high metal concentrations and low nutrients (Mormile *et al.*, 2007; Heidelberg *et al.*, 2013; Johnson *et al.,* 2015; Zaikova *et al.*, 2018). Moreover, they serve as a reservoir of novel microbial functions, such as acidophilic microorganisms that have been used for extracting metal ores from sulphide minerals. Hence, such environments are of significant interest to scientists interested in understanding the fundamental concepts of microbial mechanisms used to cope with significant stressors, as well as applied areas developing microbial consortia for bioprocessing applications (Dopson *et al.*, 2017).

Lake Magic is one of the acidic hypersaline lakes (ca. 1 km in diameter) present within the Yilgarn Craton in WA. This unique lake exhibits extremely low pH (<1.6) coupled to very high salinity (32% TDS) with the highest concentration of aluminum (1774 mg/L) and silica (510 mg/L) in the world (Bowen and Benison 2009; Conner and Benison, 2013). Lake Magic, similar to other lakes in WA, has dynamic and characteristic stages of lake transformation, including flooding, evapo-concentration and desiccation that are driven by the local seasons (Bowen and Benison 2009; Conner and Benison, 2013). The lake is fed via both regional acidic groundwater and infrequent precipitation (Benison *et al.*, 2007). Recent studies of the microbial diversity of Lake Magic has revealed that the lake hosts acidophilic, acidotolerant, halophilic and halotolerant bacterial species (Zaikova *et al.,* 2018). Moreover, fluid inclusions of halite crystals from Lake Magic, exhibited the presence of micro-algae and prokaryotes trapped within them (Conner and Benison, 2013). More recently, metagenomic analyses of lake water, groundwater and within the sediment of Lake Magic revealed that the lake is dominated by only a few species, such as *Salinisphaera*, and has a low representation of other bacterial species (Zaikova *et al.,* 2018). Interestingly, the pelagic zone of the lake was abundant in eukaryotes, including fungi and green algae, giving rise to the bright yellow colour of the lake (Zaikova *et al.,* 2018, Conner and Benison, 2013). These studies provide indicators of the population residing within the lake and the functional niches they occupy. However, they do not emphasize the survival mechanisms adopted by the resident microorganisms and how the diversity of microbial populations residing within the lake changes during different transformational stages of the lake.

Molecular approaches to understand the dynamics of Yilgarn Craton lakes in the past have focused primarily on spatial composition (Mormile *et al.*, 2009; Johnson *et al.*, 2015; Zaikova *et al.*, 2018; Aerts *et al.*, 2019). However, these studies implicitly assume that a single time point can provide a comprehensive representation of the community members. In opposition, temporal approaches to study microbial community dynamics have recently revealed considerable variability within microbial community compositions over time (Ju and Zhang, 2014; Chénard *et al.*, 2017; Nagarkar *et al.*, 2018; Cruaud *et al.*, 2019), showing that some taxa remain consistent in their abundance whilst others exhibit sudden blooms (Nagarkar *et al.,* 2018).

We hypothesize that the large fluctuations in environmental parameters during lake transformations are key drivers which will lead to marked changes in microbial populations, and contribute to the mechanisms driving the dynamics of these communities (Cruaud *et al.,* 2019). Therefore, studying the temporal dynamics of microbial communities in poly-extreme ecosystems such as Lake Magic could reveal crucial information about the trophic interactions and survival mechanisms (Nagarkar *et al.*, 2018). This study attempts to investigate the diversity and functional dynamics of bacterial, archaeal (referenced as ‘bacterial’ from hereon) and fungal communities, over a period of one year. Using 16S rRNA gene and ITS gene sequencing data, we elucidate the diversity patterns generated during lake transformations and assess likely mechanisms the resident microorganisms adopt in order to survive in the face of multiple stressors.

## 2 Materials and Methods

### 2.1 Sample collection

Samples were collected from Lake Magic for every lake stage in a span of one year (July 2017-July 2018). Multiple sampling sites were chosen around the lake to eradicate a spatial sampling effect. Sediment samples were taken as cores, which were divided into salt mat and sediment layers (**Figure 1)**. All samples were taken with sterile cores and spatulas. After every sample the cores and spatulas were sterilized with 70% ethanol. A total of 40 sediment samples and 40 salt mat samples (8 cores for each time point) were collected. Temperature, pH, and salinity were measured for each sampling point. All samples were kept frozen at −20°C in 50 ml Falcon tubes until further processing.

**Figure 1:**
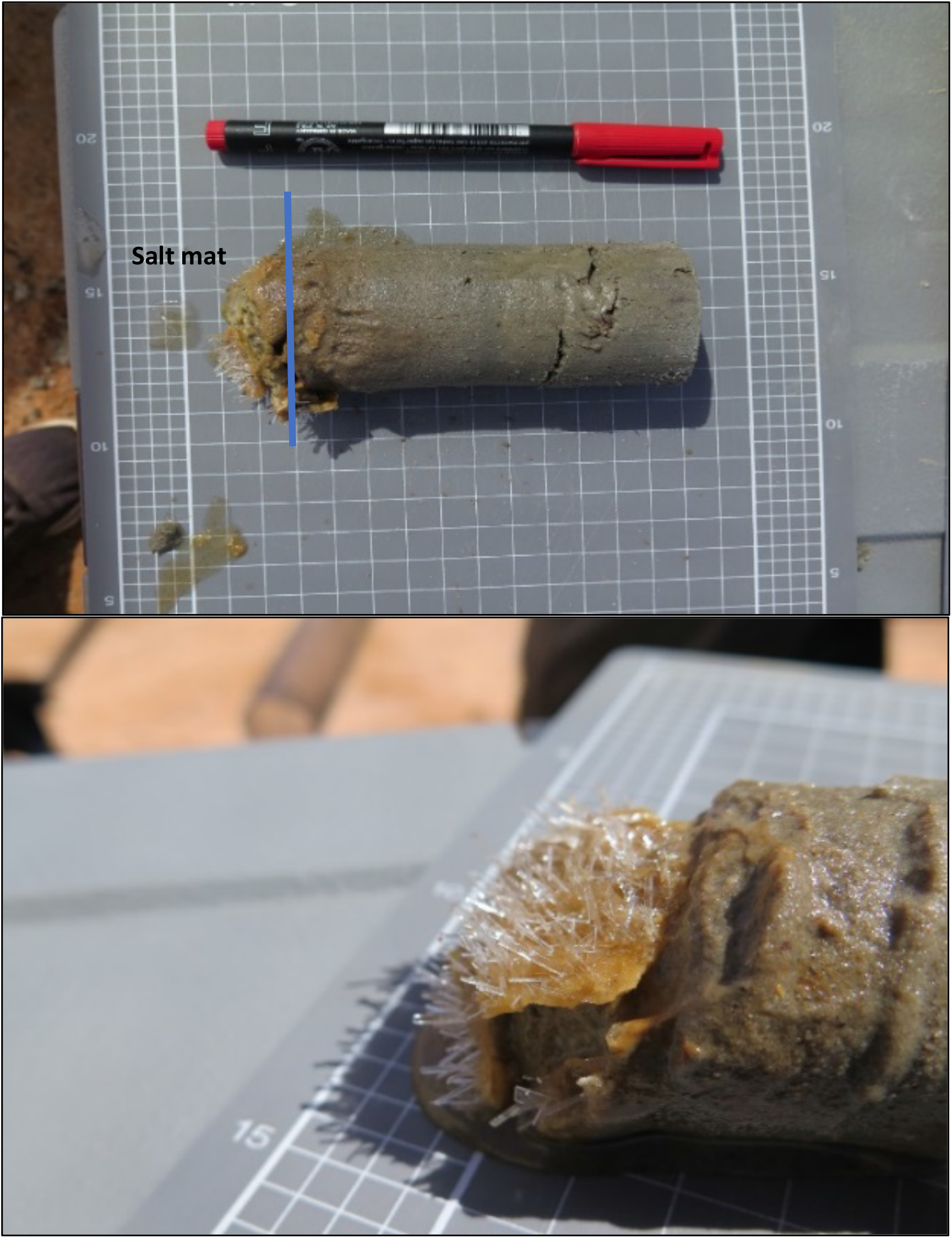
A single core of sample (~6 cm long) showing the salt mat and sediment sections. The bottom figure shows the salt mat with salt crystals on the top of sediment, sampled as a separate sample.

### 2.2 DNA extraction, PCR and amplicon sequencing

DNA from sediment and salt mat samples was extracted using a method developed for acid saline sediments. Briefly, 0.4g sediment sample in 2ml tubes containing glass beads. To this 900μl of extraction buffer (consisting of 0.2M sodium phosphate buffer, 0.2% CTAB, 0.1M NaCl and 50mM EDTA), 100 μl of 10% SDS and 10 μl of Proteinase K (20 mg/ml). The mixture was kept at −80C for 5 min and then heated at 70C for 20 min. The mixture was subjected to mechanical agitation at 20Hz for 20min after which the tubes were kept on ice for 5 min. The mixture was centrifuged at 10, 000 x g for 5 min. To the supernatant 750 μl of chilled phenol-chloroform-isoamyl (pH 8) solution was added and centrifuged for 20 min at 10, 000 x g. Aqueous layer was transferred to sterile tube and 600 μl of chilled chloroform -isoamyl (pH 8) solution was added. The mixture was centrifuged at 10, 000 x g for 5 min. Next, to the aqueous layer 650 μl of 20% PEG, 2.5 M NaCl was added and incubated at 4C for overnight. The solution was centrifuged for 20 min at 10, 000 x g. The obtained pellet was washed with 70% ice cold ethanol and 2 μl glycogen. The final pellet was dissolved in 50 μl of TE buffer. All extractions were carried out in triplicate. Tubes containing no sample were incorporated as extraction blanks (controls) and were treated identical to sample extractions. DNA concentration was measured through fluorometry using a Qubit dsDNA HS Assay Kit with a Qubit 2.0 fluorometer (Life Technologies).

Extracted DNA from all timepoints were diluted 10-fold prior to PCR amplification of the 16S rRNA and ITS genes. All PCR reactions were carried out in triplicate. PCR amplification of the 16S rRNA gene V4-V5 region was performed using the universal PCR primer set 515F and 806R, targeting members within both bacterial and archaeal domains (16). The forward primer included the addition of an Ion Torrent PGM sequencing adapter, a GT spacer and a unique Golay barcode to facilitate multiplexed sequencing. Barcoded PCR reaction mixtures (20 μl) consisted of DNA template (1 μl), universal primer mix (untagged 515F and 806R at a final concentration of 0.2μM), tagged 515F primer (0.2 μM), 600 ng BSA (Life technologies) and 2.5 x 5’Hot Master Mix (5Primer, Australia). The PCR cycle was set at 94C for 2min followed by 25 cycles of 94 min for 45 sec, 50C for 60 sec and 65C for 90. This was followed by 2 cycles of 94C for 45 sec, 65C for 90 sec and final extension at 65C for 10 min.

Amplification of the fungal component was carried out using the universal primer set ITS1 F and ITS2 R, with the addition of an Ion Torrent PGM sequencing adapter, a GT spacer and a unique Golay barcode to the forward primer. The barcoded PCR primer mixtures (20 μl) included DNA template (1 μl), universal primer mix (untagged ITS1 F and ITS2 R at a final concentration of 0.2 μM), 600 ng BSA (Life technologies), tagged ITS1 F primer (0.2 μM) and 2.5x 5’ Hot Master Mix (5Primer, Australia). The PCR conditions included initial denaturation at 94 for 2 min followed by 25 cycles of 94C for 45 sec, 50C for 60 sec and 65C for 90 sec. This was followed by 9 cycles of 94C for 45 sec, 65C for 90 sec and 65 for 10 min.

Several positive, no template (as negative controls) controls and DNA extraction controls (extraction blanks) were amplified along with the samples for both bacterial and fungal marker genes. PCR reaction performance was checked by loading PCR amplicons along with positive and negative controls on a 2% (w/v) agarose gel. The amplicons were quantified using a Qubit dsDNA HS Assay Kit on the Qubit 2.0 fluorometer (Life Technologies). All amplicons were subsequently pooled in one composite mixture at a concentration of 20 ng/μl, including negative controls. The pool was purified using AMPure XP (Beckman Coulter, Australia) and the quality of the pool was checked by visualizing on a 2% (w/v) agarose gel. The composite pool was sequenced on an Ion Torrent PGM.

### 2.3 Sequence analysis and statistical analysis

Raw sequences were de-multiplexed and quality filtered through a custom QIIME pipeline (Quantitative Insights into Microbial Ecology) (Caporaso *et al.,* 2010) with a minimum average quality score of 20. The minimum sequence length was maintained at 130 b.p. and maximum sequence length of 350 b.p. Chimeric sequences were removed using USEARCH v6.1. No forward or reverse primer mismatches or barcode errors were allowed and maximum sequence homopolymers allowed were 15. The maximum number of ambiguous bases was set at six. Denovo OTU picking was performed using ULCUST at 97% sequence identity cut off values and taxonomy was assigned through the Greengenes database (version 13.8). For fungal data, taxonomy was assigned using the SILVA v123 database (Quast *et al.*, 2012).

The OTU tables obtained for different levels of taxonomy were used as measures of taxa relative abundance in univariate statistical analysis. The OTUs detected in negative controls were manually removed from the data set. Alpha (α)- diversity at the phylum level for both 16S rRNA gene and ITS gene data was calculated using the richness, evenness and Shannon Weiner diversity index using the relative frequency table generated from a rarefied ‘biom’ table. Data normality was checked with the Shapiro-Wilk test and log transformations of the data were performed where appropriate. Differences in the diversity for each stage and layer (sediment, salt mat), was calculated using a two-way analysis of variance (ANOVA). Tukey HSD *post hoc* comparisons of groups were used to identify which groups were significantly different from each other.

Beta diversity (β) of microbial communities was calculated with nonmetric multidimensional scaling (nMDS) using Bray-Curtis dissimilarity (Bray and Curtis, 1957). Statistical significances of dissimilarity, based on temporal data and sample layer, was assessed using main effect and pairwise ANOSIM in R using the vegan package.

Phylogeny was inferred using Phylosift (Darling *et al*., 2014) for sequences that could not be classified past the domain level. The OTU abundance and diversity patterns were calculated using the vegan package (Dixon, 2003) in R software (R Core Team, 2014). Plots and heat maps were produced using ‘ggplot2’ (Wickham, 2011), ‘ggpubr’ packages in R and the online suite ‘Calypso’ (Zakrzewski *et al.*, 2017).

Predicted microbial functions of the bacteria residing within Lake Magic were generated using FAPROTAX, using default settings. FAPROTAX is a manually constructed database that maps the microbial taxa to metabolic functions (Louca *et al.,* 2016). The output from FAPROTAX was visualised in R using the ggplot2 package.

### 2.4 Chemical analysis

Ten grams of sediment sample for each time point was oven dried at 60°C until completely desiccated. The sample was crushed, sieved, packed in plastic bags and sent to the School of Agriculture and Environment (SAgE), University of Western Australia (UWA) chemical analysis laboratory. The analytical analysis included analysis of phosphorus, potassium, sulfur, organic carbon, nitrogen, iron, copper, sodium, boron, calcium, zinc, aluminum, magnesium and manganese. Concentrations were expressed as percentage per weight and mg/kg.

## 3. Results

### 3.1 Lake stages and physico-chemical parameters

A total of 5 time points representing 5 different stages of Lake Magic were sampled for this study. Although the lake did not go through some of the more extreme physical changes during the year, the transformation of one stage to the next was evident. Namely, we observed flooding, early evapo-concentration, mid evapo-concentration, late evapo-concentration and early flooding in the lake **(Figure 2)** details of which are presented in **Table 1** and **Table 2.** During the span of this study the lake did not reach complete desiccation due to heavy rainfall from 2017 to 2018.

**Figure 2:**
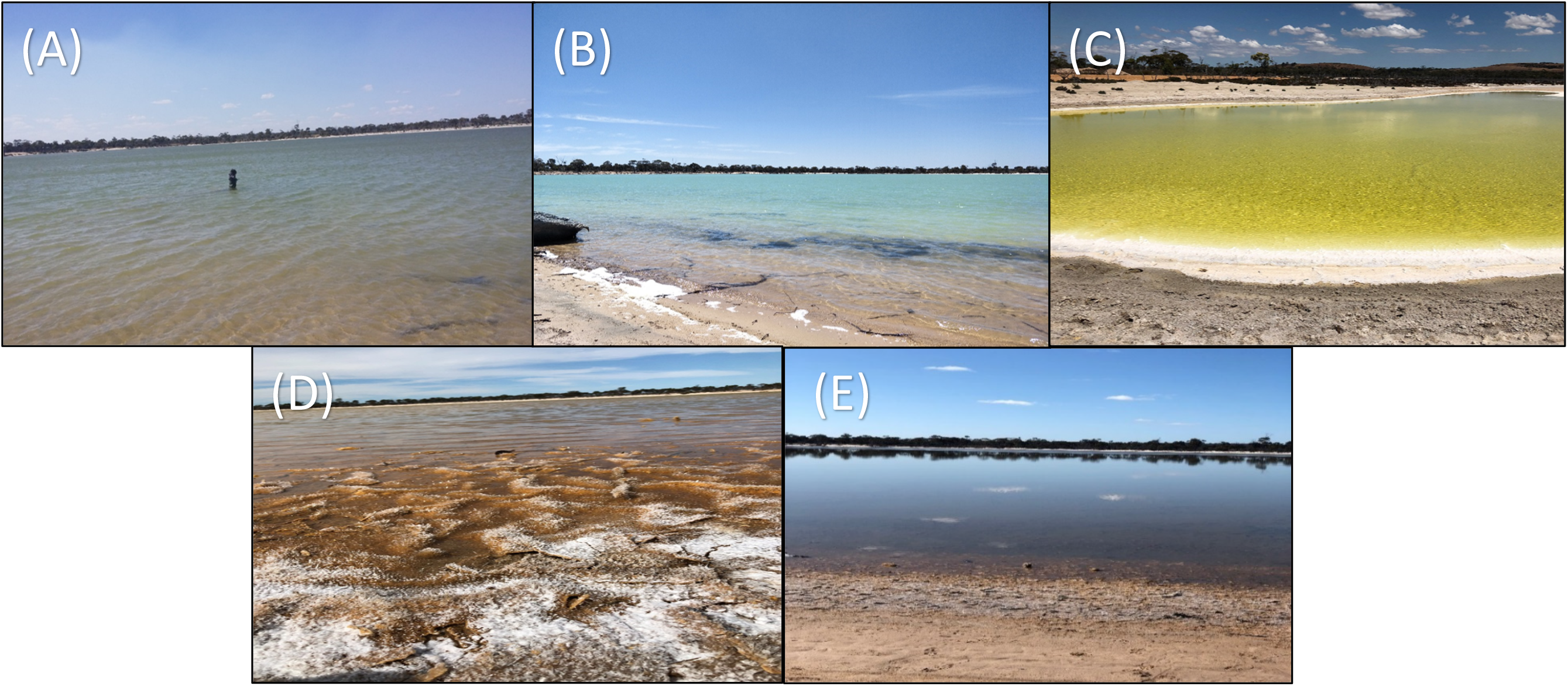
Five stages of Lake Magic in a span of one-year (a) July 2017, flooded (b) October 2017, early evapo-concentration (c) January 2018, mid evapo-concentration (d) March 2018, late evapo-concentration (e) July 2018, early flooding.

**Table 1:**
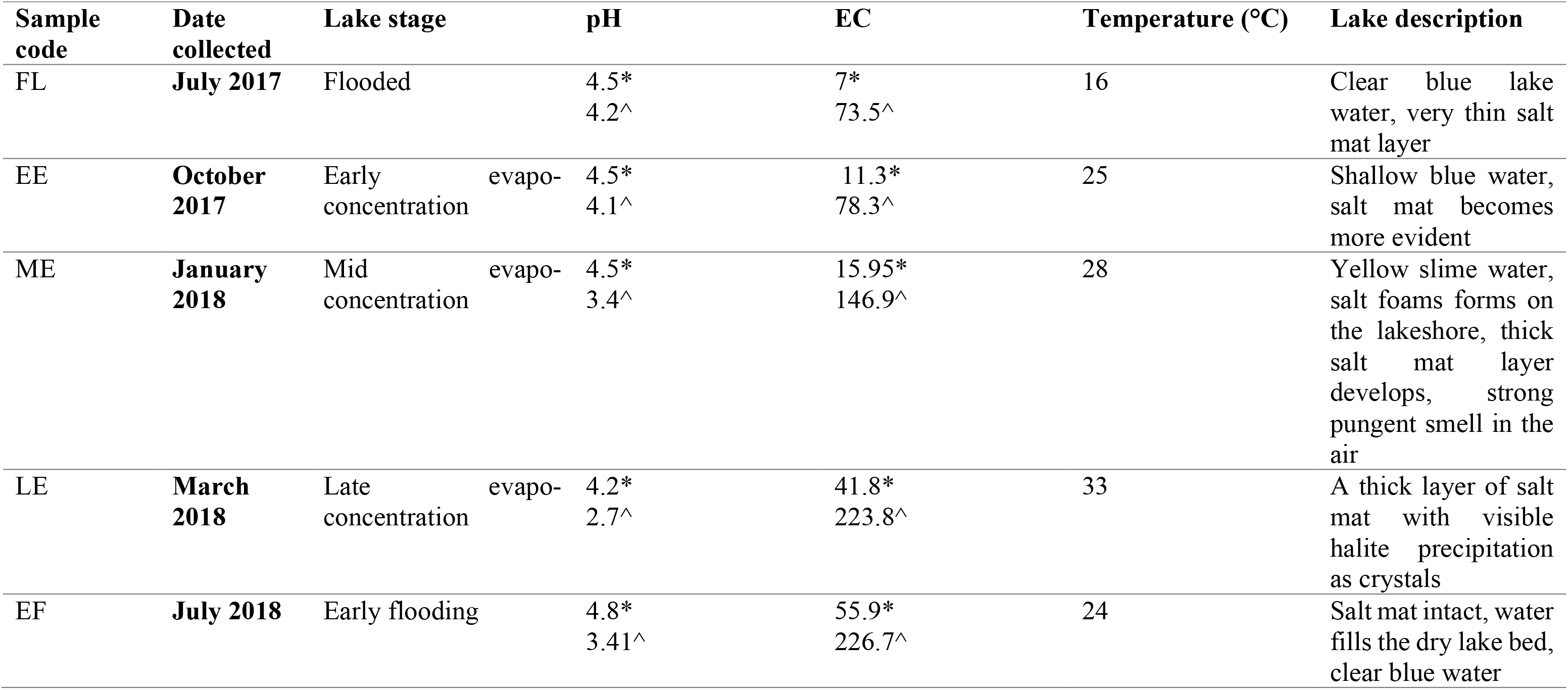
Description of sampling site at different sampling time points (* sediment, ^ lake water), (EC calculate in mS/cm)

**Table 2:** Chemical data for samples at different time points. All values shown are assessed using a single sample at each time point.

| Stage | Sampling time | % Carbon | % Nitrogen | Al (mg/kg) | B (mg/kg) | Ca (mg/kg) | Cu (mg/kg) | Fe (mg/kg) | K (mg/kg) | Mg (mg/kg) | Mn (mg/kg) | Na (mg/kg) | P (mg/kg) | S (mg/kg) | Zn (mg/kg) |
| --- | --- | --- | --- | --- | --- | --- | --- | --- | --- | --- | --- | --- | --- | --- | --- |
| Early flooding | July 2018 | 1.56 | 0.064 | 1186 | 6.8 | 10790 | 0.7 | 33 | 781 | 2661 | 1.5 | 33759 | 7.4 | 9190 | 0.2 |
| Late evapo-concentration | March 2018 | 0.739 | 0.085 | 1099 | 7.5 | 20565 | 0.6 | 77 | 1065 | 2750 | 2.5 | 37861 | 8 | 16261 | 0.2 |
| Mid evapo-concentration | Jan 2018 | 1.30 | 0.088 | 1256 | 5.7 | 2750 | 0.6 | 52 | 598 | 1462 | 1.5 | 19260 | 12 | 2591 | 0.2 |
| Early evapo-concentration | Oct 2017 | 0.597 | 0.061 | 1023 | 3.6 | 11308 | 0.4 | 47 | 357 | 733 | 0.9 | 9526 | 10 | 8716 | 0.1 |
| Flooding | July 2017 | 1.01 | 0.062 | 1007 | 3.7 | 350 | 0.8 | 64 | 323 | 587 | 0.6 | 7056 | 6 | 488 | 0.1 |

The first sampling point was during late winter to early summer in 2017, where the lake was filled with several centimeters of clear blue water (July 2017) **(Figure 2A)**. The lake sediment was rich in clay and a small amount of wet salt mat sample was acquired. In October 2017, the lake became shallower **(Figure 2B)** during early evapo-concentration **(Table 1),** characterized by clear water but where halite precipitation was evident on the lake shore. The lake transformed into a shallow yellow lake during January 2018 sampling **(Figure 2C)** where the surroundings of the lake were rich in halite precipitation, the lake itself exhibited a pungent acidic odour and the salt mat became desiccated and substantial. During mid-summer (March 2018) the lake bed became dry and a thick salt crust was observed on the surface of the sediment **(Figure 2D)** with visible salt crystals **(Figure 1),** constituting the late evapo-concentration stage. At this stage the lake was also rich in iron oxide precipitation, which was evident due to its distinct colour. The final sampling timepoint was the beginning of the flooding stage in July 2018 **(Figure 2E)** where the lake started to fill with water, with a concomitant dissolving of the halite and iron precipitation observed during the evapo-concentration stages. The chemical data has been summarized in **Table 2**.

### 3.2 Temporal dynamics of the Lake Magic microbiome

In order to assess the microbial community dynamics in the lake at different stages, we used 16S rRNA gene and ITS gene Ion Torrent sequencing (15 Salt mats; 15 Sediments) in triplicate. After DNA extraction, all samples had detectable amounts of DNA, but the DNA concentration was consistently higher for salt mat samples when compared to the sediment samples. However, there was no significant difference (ANOVA p>0.05) between diversity and richness indices of salt mat and sediment layers for both bacteria and fungi analyses **(Supplementary Figure 1),** likely indicating minimal diversity analysis bias despite wide variations in DNA extraction concentrations. Generally, we found DNA extraction easier from salt mat samples, relative to sediment samples, due to the lower amount of clay present in the salt mat samples.

A total of 330,913 reads were obtained for 16S rRNA microbiome analyses, representing 15947 OTUs. The OTUs were assigned to 37 phyla, 135 classes, 260 orders, 426 families and 739 genera of archaea and bacteria. Only OTUs which appeared in sequenced negative controls were discounted from the analyses and no other OTUs were filtered. This is because low abundance OTUs can have a significant effect on the diversity metrics for the microbial communities in low biomass environments, hence, filtering of OTUs with low representation in the microbiome can result in loss of crucial information. Analysis of the ITS gene sequences revealed a total of 724,490 reads, representing 4005 OTUs and were assigned to 15 phyla, 44 classes, 86 order, 177 family and 259 genera.

### 3.3 Bacterial communities are dynamic in Lake Magic

The Lake Magic microbiome OTU richness was analyzed for salt mat and sediment samples **(Figure 3A)** at phylum level which varied significantly between the five stages of the lake (ANOVA p= 0.001). The richness index varied from 9 to 23 for 16S rRNA gene and the OTU richness was significantly higher during the EF stage (mean richness = 21). The second highest OTU richness was observed during the ME stage (mean richness = 18). The LE stage showed the lowest mean richness of 13.6, whereas, OTU richness at the FL stage was significantly different and showed a mean richness of 16. A mean richness of 15.5 was seen during EE stage.

**Figure 3:**
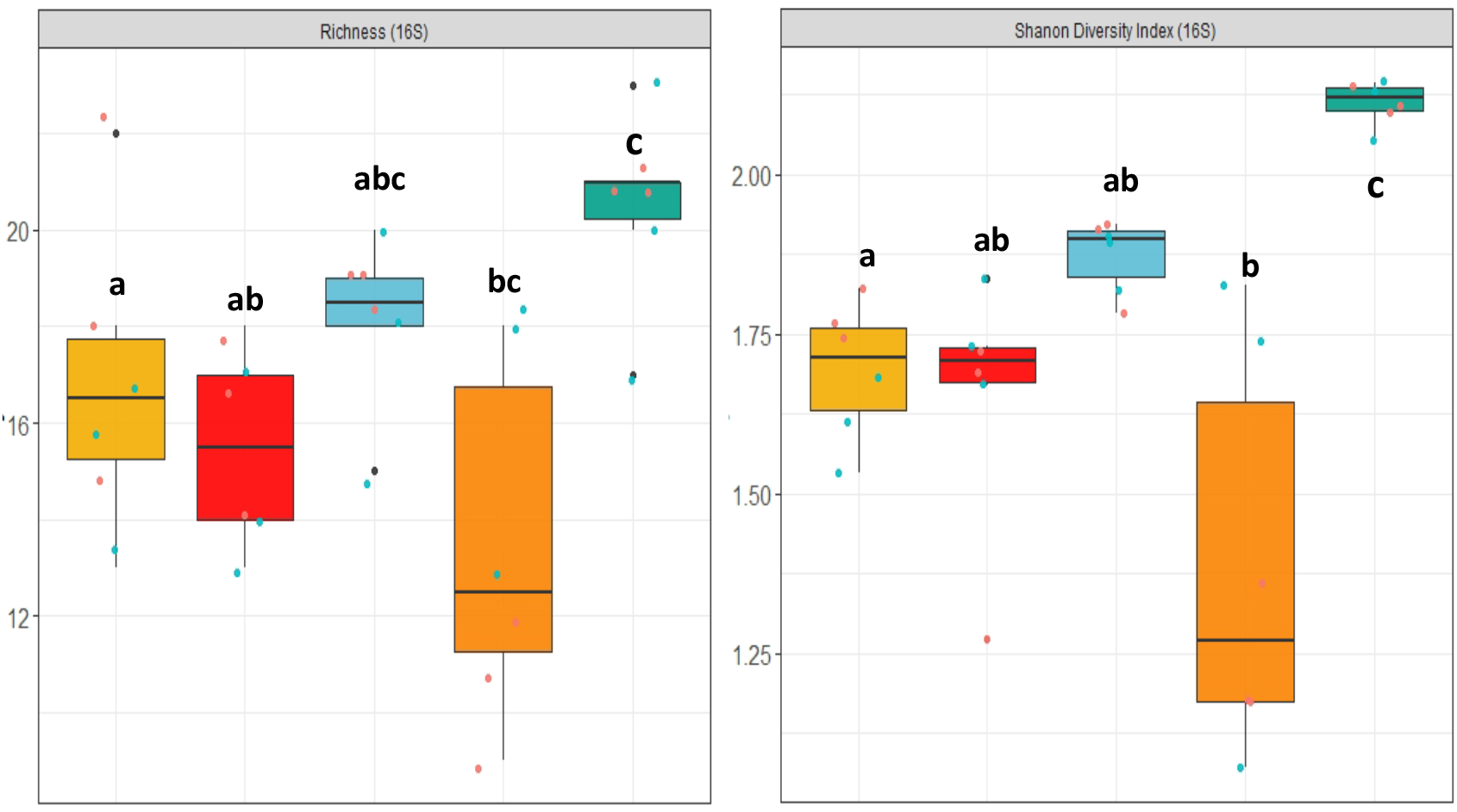
(a) 16S rRNA gene OTU richness (ANOVA, p= 0.001) and (b) Shannon diversity index (ANOVA, p<0.001) at phylum level. The letters show significant differences as obtained with Tuckey post hoc test. Colours represent: 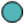 Sediment 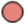Salt mat.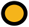FL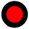 EE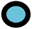 ME.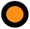LE 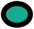 EF

The Lake Magic microbiome diversity varied during the different lake stages **(Figure 3B)** and followed a similar trend to that of OTU richness. The Shannon diversity index ranged from 1.61 to 2.13 for bacterial communities, where the diversity was seen to increase during EE (mean diversity = 1.65) and the ME stage (mean diversity =1.87). However, the diversity decreased during the LE stage and was recorded as the lowest diversity index (mean diversity= 1.39) of all stages. The highest diversity was seen during the EF stage (mean diversity= 2.11). The ANOVA test was further analyzed with a *Tuckey post hoc* test for diversity and richness indices, which revealed that the diversity during FL, LE and EF stages were significantly different from each other, whereas, the richness was significantly different only during FL and EF stages.

The Lake Magic bacterial composition under different lake stages was analyzed and are shown in (**Supplementary Figure 2).** Archaeal community diversity in the microbiome was represented by only two phyla, the *Crenarchaeota* and the *Euryarchaeota,* whilst the bacterial domain contributed the dominant microbial sequences observed within the 16S rRNA gene analyse**s.** The majority of the sequences for bacteria originated from two phyla: *Bacteriodetes* (20%) and *Proteobacteria* (39%), most of which could not be classified below family level, indicating that a large proportion of the bacterial taxa in Lake Magic appear to be relatively poorly characterized.

### 3.4 The bacterial community becomes more specialized as stress increases

For taxa which could be classified to genera, varying trends were observed during different lake stages and within the sample layers **(Supplementary Figure 3)**. Specifically, key bacterial genera fluctuated within the salt mat and sediment during the various lake stages. The correlation analysis of chemical data with the bacterial diversity revealed that microbial dynamics is strongly driven by salinity, temperature, pH and carbon content in the lake **(Supplementary Figure 4)**. For instance, members of the *Acidiphilium* genus were low in abundance during the FL stage in both the salt mat and sediment samples, whilst their relative abundance was seen to increase within in the salt mat during evapo-concentration stage as the lake conditions became more stressful. A significant increase (ANOVA, p<0.05) in *Acidiphilium* relative abundance was observed within the salt mat during the LE stage and significantly decreased during the EF stage of the lake **(Supplementary Figure 3,***Acidiphilium*). Similarly, the *Acidobacterium* genus’ relative abundance was high within sediment during the FL stage and a significant increase in abundance within the salt mat was seen during all evapo-concentration stages **(Supplementary Figure 3,***Acidobacterium*). Similar to the *Acidiphilium, Acidobacterium* abundance decreased when the lake was in the EF stage. In contrast, sequences belonging to the *Arthrobacter, Bacillus, Flavbacterium and Sporosarcina* genera increased significantly in abundance during the EF stage in both the sediment and salt mat, whilst genera such as *Nitrososphaera* were only present during the EF stage in high abundance.

Interestingly, it was also observed that the abundance of the *Sulfurimonas, Syntrophobacter, Halothiobacillus, Acidobacterium, Acidiphilium and Alicyclobacillus* genera decreased during the EF stage in both the salt mat and the sediment whilst, the *Syntrophobacter* population increased in relative abundance within the sediment during LE. During the FL stage *Salinisphaera* was found to be more abundant in the salt mat when compared to the sediment. However, when compared to the LE, its abundance increased in the salt mat **(Supplementary Figure 3).** Overall, from these data it was evident that the sediment becomes more dominated by bacteria that were not specialized for surviving in the environmental conditions, whereas, the salt mat became dominated with specialized microbes such as *Salinisphaera* and *Acidobacterium*.

### 3.5 The fungal community becomes less diverse under extreme conditions within Lake Magic

The diversity and richness of the fungal community within Lake Magic significantly decreased (ANOVA, p < 0.001) as the lake conditions became more extreme. The richness index ranged from 2 to 7 whereas the Shannon diversity index ranged from 0.004 to 1.28 **(Figure 4A and 4B).** The highest mean richness was observed for the FL and EE stage. The OTU richness consistently decreased after the EE stage, with lowest OTU richness observed for the EF stage (mean richness = 2). The fungal diversity showed a varying trend when compared to OTU richness, where the highest diversity index was observed for the FL stage (mean diversity index =1.11) whereas the lowest was observed for the extreme EF stage (mean diversity index= 0.28). A *Tuckey post hoc* test revealed that the FL and EE stage were statistically similar to each other, whilst, the ME and LE stages were statistically similar, and the EF stage was significantly different from all other stages. Additionally, a *Tuckey* test for OTU richness showed that the FL, EE and EF stages were significantly different from all other stages.

**Figure 4:**
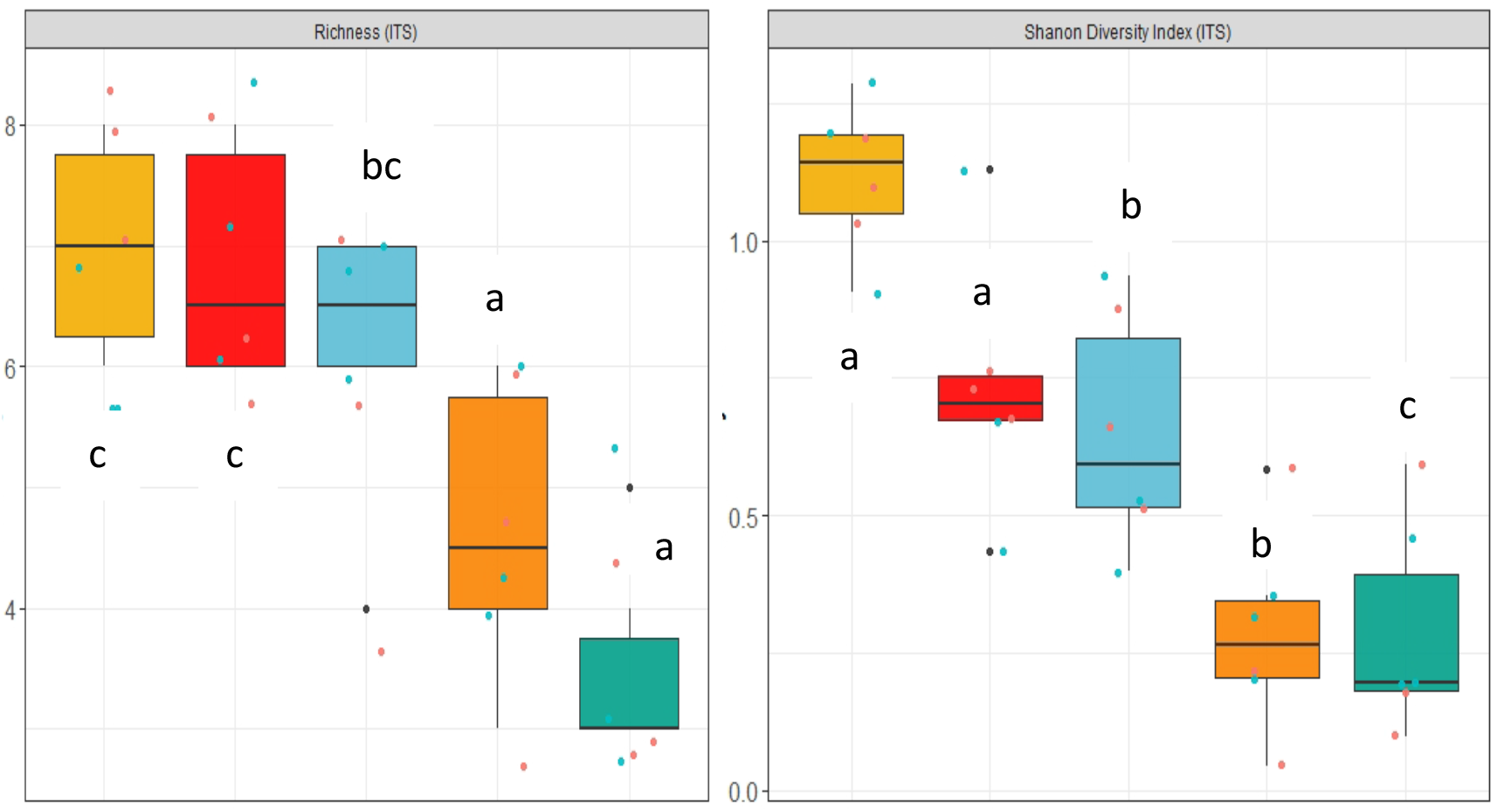
**(a)** ITS gene OTU richness (ANOVA, p= 0.001) and (b) Shannon diversity index (ANOVA, p<0.001) at phylum level. The letters show significant differences as obtained with *Tuckey post hoc* test. Colours represent: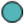 Sediment 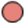 Salt mat 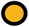FL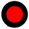EE 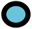ME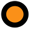 LE 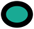EF

Interestingly, when the fungal community composition was visualized **(Supplementary Figure 5)**, an increase in unidentified fungi belonging to the Ascomycota phylum was observed. This indicated that a large portion of the fungi living in Lake Magic are likely unidentified. The salt mat during the EF stage was, however, seen to be the most diverse when compared to other time points and was abundant with the *Cladosporium* genus. Variation in the composition of other genera including *Fusarium, Ulocladium* and *Hostaea* was also seen.

### 3.6 Beta diversity of the bacterial and fungal communities

The dissimilarity between microbial communities at different lake stages was assessed using Nonmetric multidimensional scaling (nMDS) and Bray-Curtis dissimilarity indices. The sediment and salt mat bacterial communities from the FL, ME, LE and EF stages tightly clustered together **(Figure 5A)**. In contrast, the salt mat and sediment communities under the EE stage clustered separately (ANOSIM, R^2^= 0.49, p=0.001). However, similar to alpha diversity, when an ANOSIM test was applied to determine the variability in the salt mat and sediment samples no significant difference was observed.

**Figure 5:**
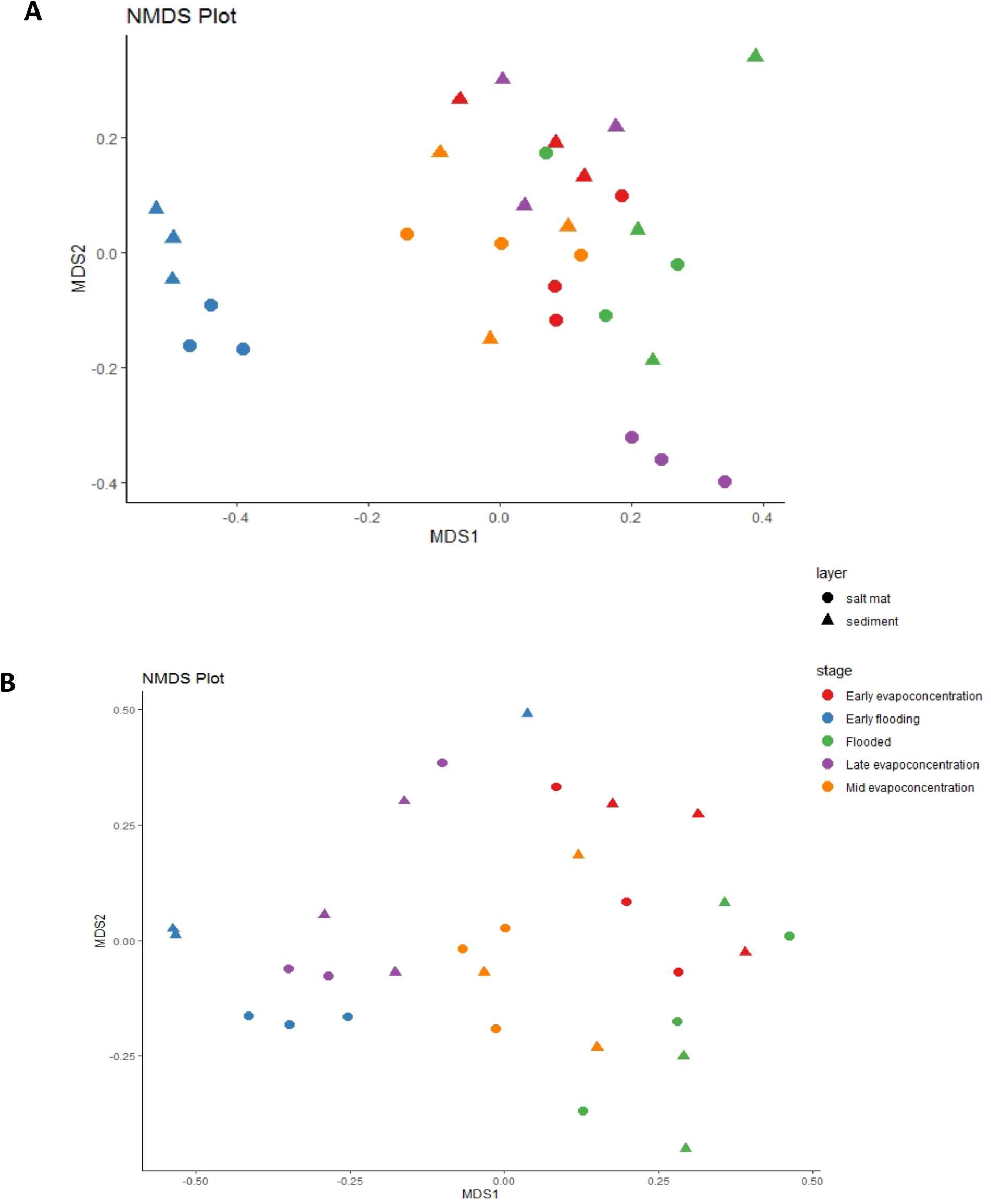
Beta diversity of (A) 16S rRNA (B) ITS sequences for different time points and layers. Samples are coloured according to the time points and shaped according to the layer

The nMDS analysis of fungal communities showed less clustering when compared to the bacterial community analysis **(Figure 5B),** where none of the lake stages clustered separately. This indicated that the members identified within the *Ascomycota* phylum vary for different lake stages (ANOSIM, R^2^ =0.462, p= 0.001), but no significant difference was seen between sediments or salt mat samples.

### 3.7 Predicted ecological functions of the bacterial communities

The ecological function of the bacterial communities analyzed was predicted using the FAPROTAX pipeline for the different stages of the lake. These analyses indicated aerobic chemoheterotrophy, iron oxidation and sulphur related pathways as likely dominant functions within the lake’s bacterial community **(Figure 6)**. Functions related to carbon metabolism, such as methylotrophy and methanotrophy, were predicted but were not abundant. Nitrogen related functions included nitrification and ammonia oxidation and were observed to fluctuate between different lake stages. For instance, these data indicated that nitrification and ammonia oxidation were significantly higher within the salt mat during the EF stage **(Supplementary Figure 6)**. Interestingly, the nitrogen related activity was also high in the sediment samples during the EE and ME lake stage. Sulphur related functions were lower during the EF stage, but significantly increased in the sediment layer during all evapo-concentration stages. Sulphur related activity was found to be highest in the sediment during the EE stage. Iron oxidation and reduction was relatively low at all timepoints, but significantly increased during the EE and LE stage within the sediment layer. Finally, Iron based respiration pathways fluctuated more frequently when compared to sulphur and nitrogen related functions during all lake stages **(Supplementary Figure 6)**.

**Figure 6:** Dot plot showing the FAPROTAX predicted functions of the bacterial community within the salt mat and sediment samples at different lake stages.

## 4. Discussion

Acid saline lakes in WA host unique microorganisms that can be studied to understand life under extreme conditions, novel biogeochemical processes and new biotechnological avenues (Benison *et al*., 2007). In this chapter, the bacterial and fungal microbial community dynamics were studied using a temporal approach to resolve how an extreme lake microbiome changes during different stress phases due to changes in the physico-chemical properties of the habitat.

Previous studies on acidic hypersaline lakes in WA revealed a high level of bacterial diversity within Lake Magic and other WA lakes water (Mormile *et al.,* 2009; Zaikova *et al.*, 2018), but have relied upon single time (lake stage) sampling points. Our results indicate that the alpha (α)-diversity of the bacterial and fungal microbial communities differed significantly between the less extreme (FL) stage and the more extreme (LE) stage. These differences indicated that the lake microbiome diversity is driven to a large degree by the high salt and low pH conditions within the lake (Podell *et al.,* 2014). It has previously been reported for hypersaline environments that the ionic concentration in these environments hinders the solubility of oxygen, and hence, the oxygen concentration is very low in acid saline lakes (Sherwood *et al.,* 1991). However, the majority of microorganisms isolated from hypersaline environments are aerobic heterotrophs who are capable of anaerobic facultative fermentation (Dyall-smith, 2009). Our results are also in line with these previous findings. Moreover, our results show that the number of OTUs in the salt mat layer were consistently higher in all samples. These results are also similar to the results obtained by Aerts *et al.*, (2019), for four acid saline lakes in Western Australia, and suggest that the majority of the microbial community diversity resides in the salt mat, where it likely has increased access to light, water and oxygen. This is likely explained by the availability of oxygen at the air/water interface, which exhibits an unequal distribution of oxygen between the salt mat and the sediment layer. Since the salt mat is directly in contact with the water column in the lake there is higher availability of oxygen locally (Podell *et al.,* 2014). Comparing the microbial composition of the salt mat and sediments in Lake Magic also indicates that microorganisms more tolerant to high salt and pH conditions are selected within the salt mat, where the conditions are harsher than those in the sediment.

The prokaryotic community at all stages of the lake cycle was dominated by halotolerant and acidophilic microorganisms, whilst archaeal taxa were found in much lower abundances. Archaeal sequences were derived from only two phyla and these observations are in good agreement with Johnson *et al.,* (2015) who observed low archaeal diversity within four acidic hypersaline lakes in Yilgarn Craton. It has been reported for Lake Tyrell in Victoria, Australia, that succession of different microorganisms is dependent upon the solutes present in the lakes (Podell *et al.,* 2014). Hence, the variation in solute concentration in Lake Magic at different stages is likely driving the succession of archaea and bacteria. It could also be hypothesised that bacterial taxa outcompete archaeal taxa in these unique environments because of the dynamic nature of the WA lakes, where environmental changes are frequent and, therefore, stress tolerant species are selected, as opposed to obligate archaeal extremophiles (Benison and Bowen, 2006).

Organic carbon and nitrogen values were found to be low at all lake stages and increased slightly during the flooded (FL) stages, in line with previous studies in similar lakes in WA (Ruecker *et al.,* 2016; Aerts *et al.*, 2019). Similarly, phosphorus levels were also low in Lake Magic, as in other WA lakes (Aerts *et al.*, 2019), with phosphorus often being considered a limiting factor along with carbon and nitrogen for the growth of microorganisms (Elser *et al.,* 2007). Microbiome community analyses indicated an abundant community of heterotrophs within the lake and hence, low carbon, nitrogen and phosphorus are likely to be responsible for the low the biomass in these samples (Aerts *et al.*, 2019). However, carbon and nitrogen levels were highest during the ME and these increases are likely due to photosynthetic inputs via blooming of the saline tolerant algae *Dunaliella.* This is also reflected in the increased microbial diversity during the ME stage. Blooming of algae commonly occurs during this lake stage and provides a source of photosynthetically derived nutrient input which causes increased diversity in the lake (Zaikova *et al*., 2018).

During the evapo-concentration stages iron concentrations increased which was also mirrored in the predicted microbiome functions of increased iron respiration activity within the bacterial community. Iron metabolism is considered an important function of the sediment inhabitants and is thought to be responsible for the lowering of pH in the sediment (Lu *et al.*, 2016; Zaikova *et al.*, 2018). The *Alicyclobacillus* genus members are reported to be involved in iron oxidation along with archaea in other acid saline lakes in WA (Johnson *et al.*, 2015; Lu *et al.*, 2016). The members of the genus are also capable of reducing iron (Yahya *et al.,* 2008; Lu *et al.*, 2010). Previous metagenomic studies of Lake magic indicated that *Alicyclobacillus* was the most abundant genus within the sediment. Moreover, the *Acidiphilium* genus was also found to be abundant in the sediment samples (Zaikova *et al.,* 2018) with members of this genus are involved in iron reduction (Weber *et al.,* 2006; Sánchez-Andrea *et al.*, 2011; Zaikova *et al.*, 2018).

These data indicated, that the abundance of the *Alicyclobacillus* and *Acidiphilium* genera varied across different lake stages and is likely more complicated than first thought, in terms of lake biogeochemistry (Zaikova *et al.,* 2018). *Alicyclobacillus* abundance was significantly higher in sediment samples during the FL stage and was significantly increased during the EE stage within the salt mat samples. In contrast the, *Acidiphilium* genus was more abundant within the salt mat samples during the later LE stage. Interestingly the overall abundance of these genera decreased during the higher stress stages of the lake. Previously it has been suggested that archaea dominate iron oxidation in acid saline extreme environment and that the role of bacteria is limited. Moreover, the Iron activity is known to decrease by the presence of high salt in the environment (Lu *et al.,* 2016). In the study, *Acidiphilium* spp. were isolated from an acid saline lake in WA which formed long filamentous structures during low pH and high salinity conditions indicating that these species have developed a coping mechanism for extreme stress (Lu *et al.,* 2016). Examining the functional profile of our samples it can be seen that predicted iron respiration would increase significantly in the sediment during the EE and LE stage and is the lowest in the sediment during the FL stage potentially because of the coping mechanism adapted by these species.

The activity of acidophiles is known to decrease significantly in presence of high concentration of chloride ions (Suzuki *et al.*, 1999; Shiers *et al.*, 2005). However, Lu *et al.,* (2016) found that *Acidiphilium* spp. formed long filaments as a coping mechanism to high chloride ions within the Dalyup river in WA. These data, in concert with previous results, suggest that iron metabolism is affected by the presence of chloride ions and we hypothesise that this may explain the bacterial spp. of *Alicyclobacillus* and *Acidiphilium* being dominant during EE and FL stages. These data also suggest that under the high chloride conditions, based upon community data, these species are responsible of lowering the pH in the sediment. Hence, we postulate from these results that microorganisms in Lake Magic adapt to stressful conditions of pH and salinity not only physiologically, but also play crucial roles in changing their external environment making it more habitable for themselves and other members of the Lake.

One of the most abundant genera detected in these data was the *Salinisphaera* (7%), in both the sediment and salt mat. *Salinisphaera* species are mesophilic, halotolerant and slightly acidophilic (pH range 5.0-7.5), surviving in a range of moderately acidic and saline conditions as well as high concentrations of metal ions. Species belonging to the *Salinisphaera* genus have been isolated from a range of environments, including hydrothermal vents, solar salterns, brine from salt wells, seawater and marine fish surfaces (Antunes *et al.*, 2003; Mormile *et al.*, 2007; Crespo-Medina *et al.*, 2009; Bae *et al.*, 2010; Gi *et al.*, 2010; Park *et al.*, 2012; Zhang *et al.*, 2012; Shimane *et al.*, 2013). Critically, these species are able to metabolize both autotrophically and heterotrophically (Antunes *et al.,* 2003; Crespo-Medina *et al.*, 2009; Bae *et al.*, 2010; Antunes *et al.*, 2011a; Park *et al.*, 2012; Zhang *et al.*, 2012; Shimane *et al.,* 2013). In addition, *Salinisphaera* species are also involved in the uptake of iron and siderophore production (Antunes *et al.*, 2003). Molecular studies here indicated that the abundance of *Salinisphaera* was consistently high in salt mats during all lake stages, except during the flooded (FL) stage, where it was more abundant within the sediment. *Salinisphaera* was previously reported to be the single most dominant OTU in the lake water (Zaikova *et al*., 2018) during evapo-concentration in Lake Magic. These findings together indicate that most of the *Salinisphaera* spp. reside in the water column of Lake Magic. The dramatic increase in its representation during the LE stage in the salt mat and sediment suggests that it is highly tolerant of extreme pH and acidic environmental conditions, including tolerance to heavy metal ions, allowing it to survive through the evapo-concentration stages.

Members of the *Sulfurimonas* genus have been isolated from diverse environments such as hydrothermal vents, marine sediments and terrestrial habitats and are known to play an important role in chemoautotrophic processes (Han and Perner, 2015). The members of this genus can grow on a variety of electron donors and acceptors and, thus, are able to colonize disparate environments. These include different reduced sulphur compounds such as sulphide, elemental sulphur, sulphite and thiosulfate (Han and Perner, 2015). Many members of the genus are also involved in nitrogen and hydrogen metabolism. In our samples, *Sulfurimonas* (1.2%) abundance continuously decreased in salt mats but increased in abundance within the sediments as the environmental conditions became more stressful. Ultimately, it decreased significantly during the most extreme (LE) lake stages but still maintained a low representation in the sediment. When examining the functional profile data, sulphur respiration significantly increased in the sediment samples during the evapo-concentration stages, suggesting that *Sulfurimonas* spp play a crucial role in cycling key nutrients in the sediment. Since the members of the genus are able to survive chemolithoautotrophically using various electron acceptors and donors (Campbell *et al.,* 2006; Grote *et al.,* 2008), it suggests that these species are capable of adapting to the changing environmental conditions of Lake Magic. A similar trend of abundance was seen for *Flavobacterium, Bacillus and Syntrophobacter* genera where their abundances increased in the sediment during the evapo-concentration stages and we postulate that they play a crucial role in niche construction by interacting with other dominant taxa as the stress increases.

The fungal diversity was seen to decrease as the environmental conditions became more extreme. A single phylum, Ascomycota (76.8%) became dominant as the lake became dry and hypersaline. Most of the members of this phylum are unidentified, and hence, most of the fungi residing in Lake Magic are either novel, or the sequence length of the marker gene used in this study is not sufficient to be able to characterise these members. It can be deduced that these fungi are tolerant of the acidic hypersaline conditions of the lake. In a previous study on Lake Magic microbiology, the water samples were found to be highly diverse in terms of eukaryotic community composition, being abundant (~98.5%) in Ascomycota. *Aspergillus* and *Penicillium* were the most abundant genera in Lake Magic water. However, the results for sediment samples were quite similar to our results (Zaikova *et al.*, 2018). The presence of the halotolerant algae *Dunaleilla* during evapo-concentration stages is a major contributor to carbon content in the lake. It can be deduced from the data that as *Dunaleilla* increase in the Lake (during ME) the diversity in the salt mat increases rapidly. Interestingly, this difference in the diversity of lake water and sediment column indicates the exchange between the two compartments in terms of nutrition. The water column is more abundant in oxygen and has access to sunlight whilst, the higher composition of eukaryotes in the water column which contribute to the carbon and nitrogen content in the salt mat and the sediment.

In conclusion, our results highlight that the diversity in Lake Magic is strongly driven by the pH and salt concentrations at different stages of the lake. In both the salt mat and sediment samples, bacteria were found to be more abundant in the lake, comprising of halotolerant and acidotolerant species and only a small representation of archaea was seen. We can conclude from our results that, the sediment was seen to become more specialised in microorganisms involved in buffering their external environment, evident from the increase in abundance of acidotolerant and halotolerant species involved in various functions such as sulphur and iron metabolism. Moreover, due to microbial activity the environmental conditions do not change to the same degree in the sediment when compared to the salt mat, resulting in a safe haven for microbes, where they are able to thrive during extreme conditions. These findings confirm our hypothesis that during stressed conditions, microorganisms extensively rely on associations with other microorganisms and the interactions increasingly become positive (Piccardi *et al.,* 2019). This also suggests that microbial interactions are involved in stabilising the ecosystem and is responsible for the resistance and resilience of these communities (Zengler and Zaramela, 2018; Naidoo *et al.*, 2019).

