## Supplementary figures for "Temporal Microbial Community Dynamics within a Unique Acid Saline Lake"

### Supplementary File

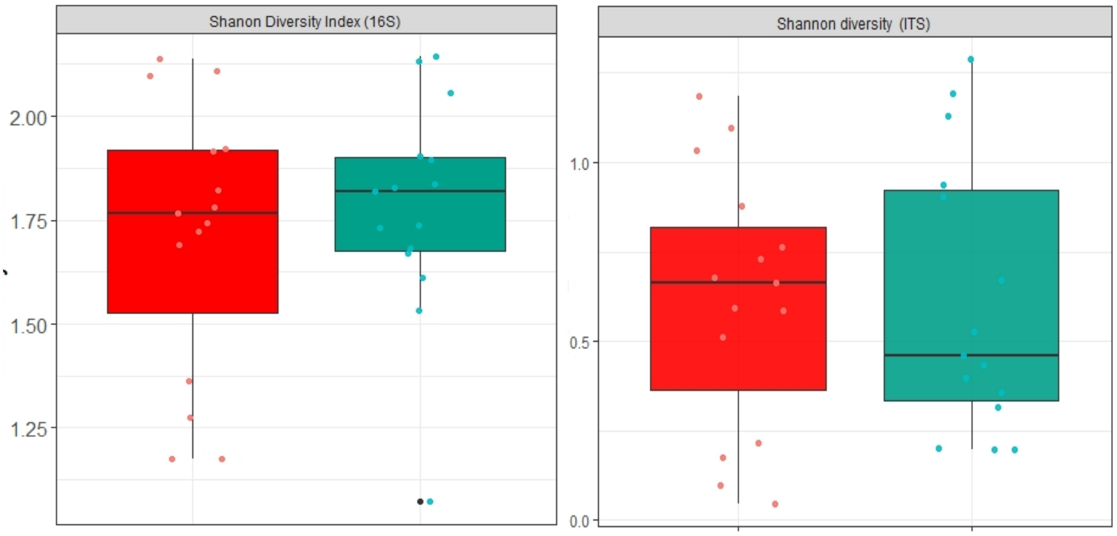


**Supplementary Figure 1:** Shannon diversity index of 16S rRNA gene and ITS gene between salt mat and sediment layers at phylum level. No significant difference was seen for diversity between the layers (ANOVA, p>0.05). Colours represents: Salt mat Sediment

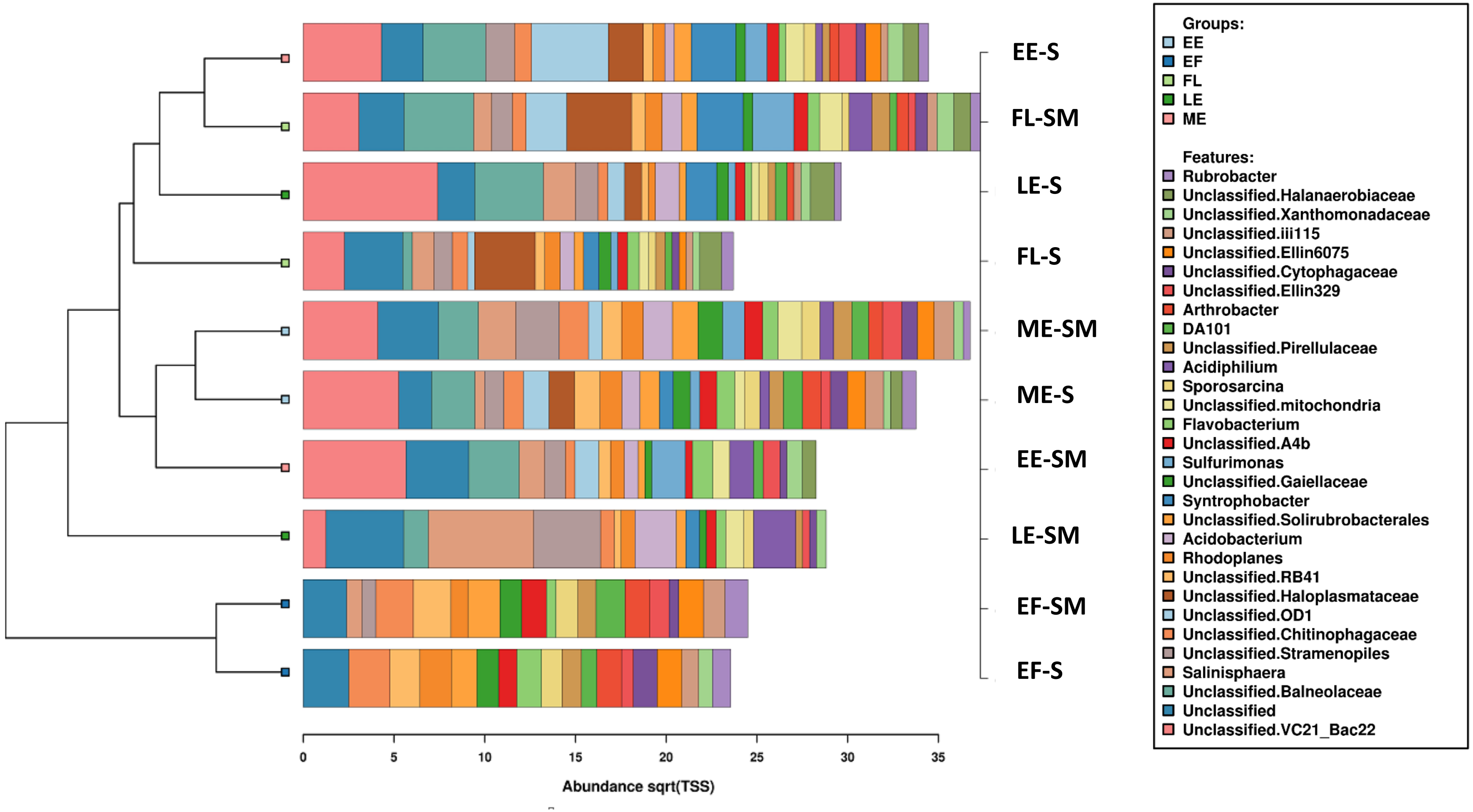


**Supplementary Figure 2:** Bar chart showing bacterial composition at genera level for each sample layer at different time points. The samples have been arranged so more closely related are clustered together.

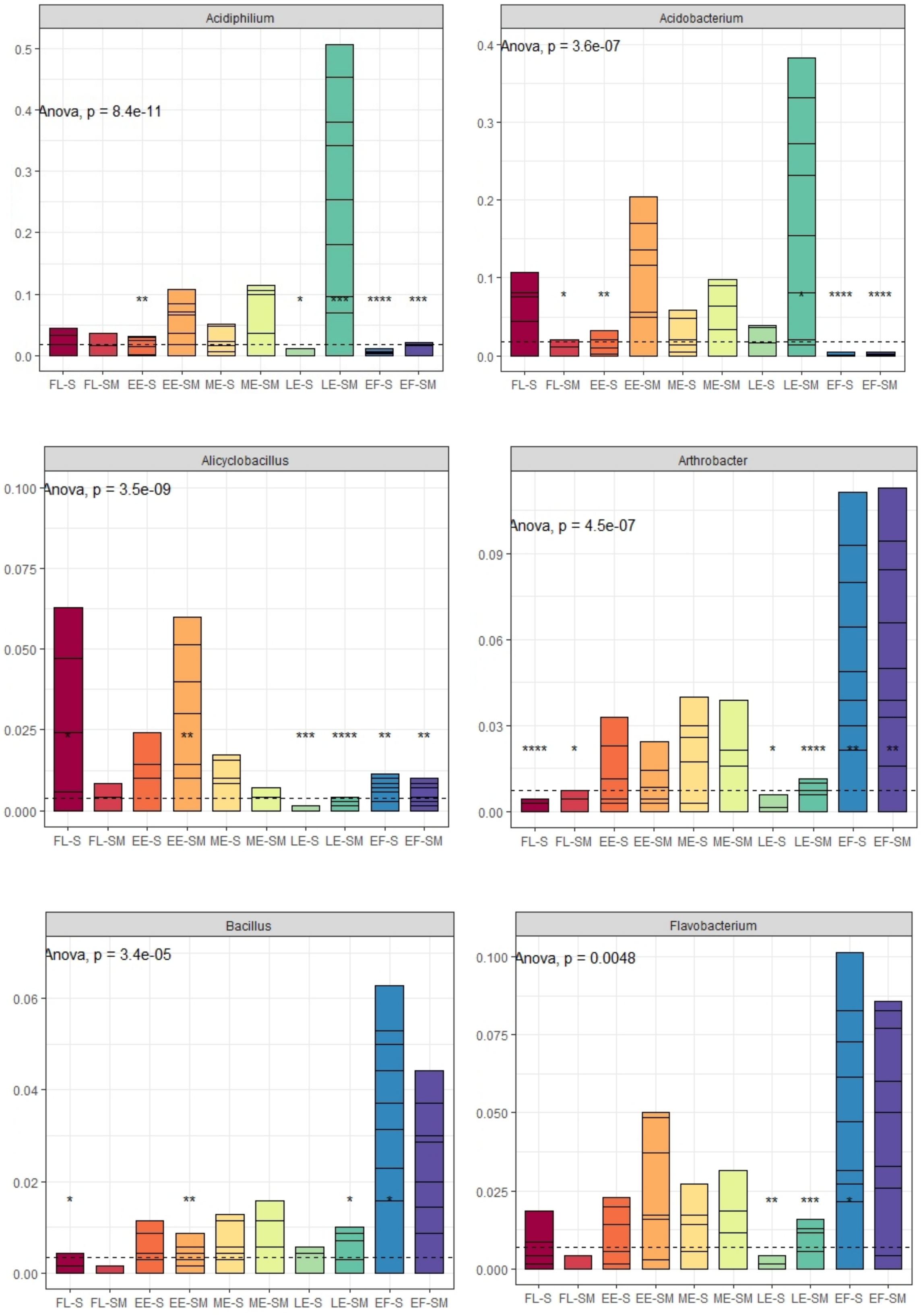
Acidiphilium
Anova, p = 8.4e-11
Acidobacterium
Anova, p = 3.6e-07
Alicyclobacillus
Anova, p = 3.5e-09
Arthrobacter
Anova, p = 4.5e-07
Bacillus
Anova, p = 3.4e-05
Flavobacterium
Anova, p = 0.0048
FL-S FL-SM EE-S EE-SM ME-S ME-SM LE-S LE-SM EF-S EF-SM

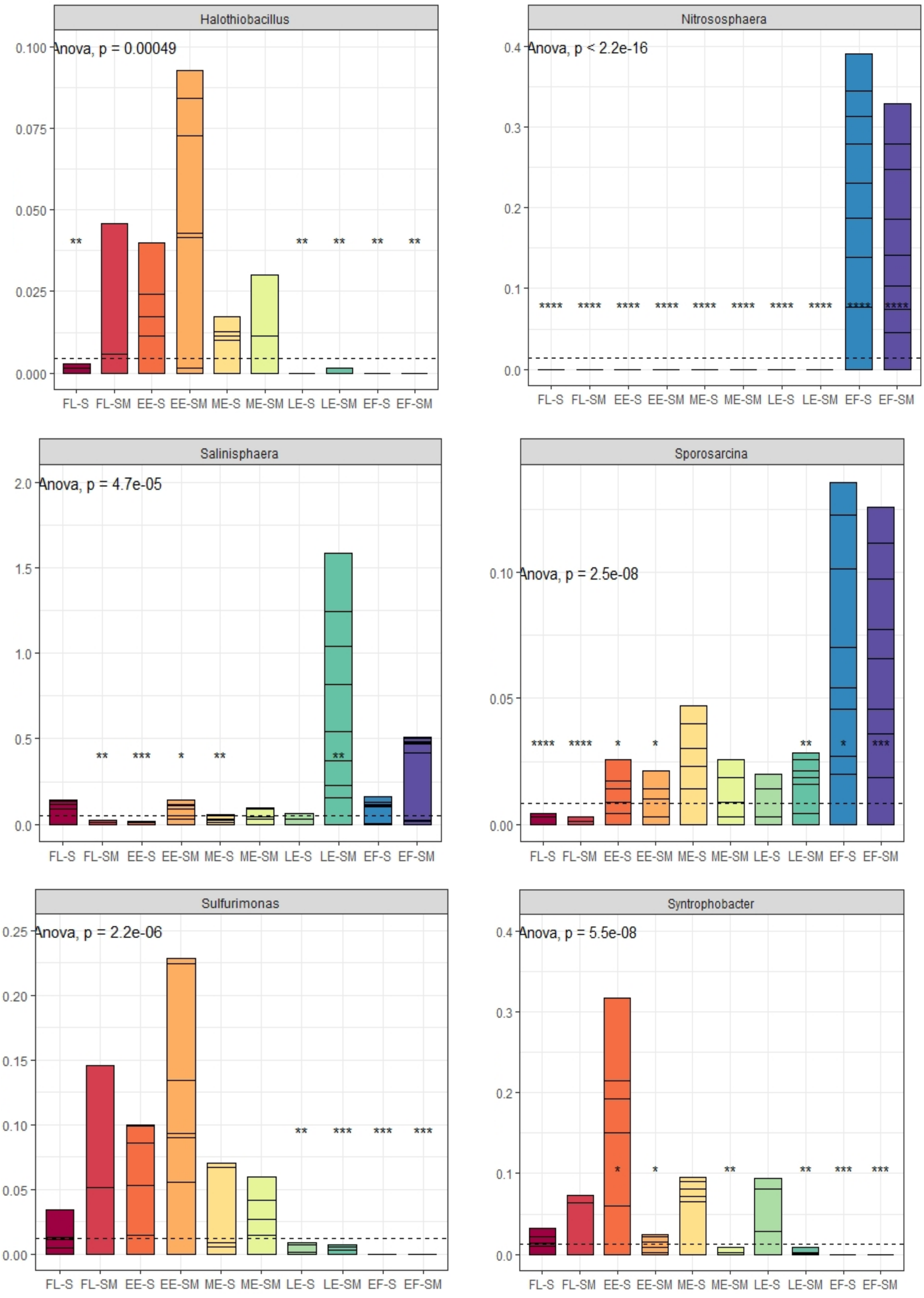


**Supplementary Figure 3:** Abundance of key genera during different stages and sample layers. Statistical significance is shown as asterisks (ANOVA).

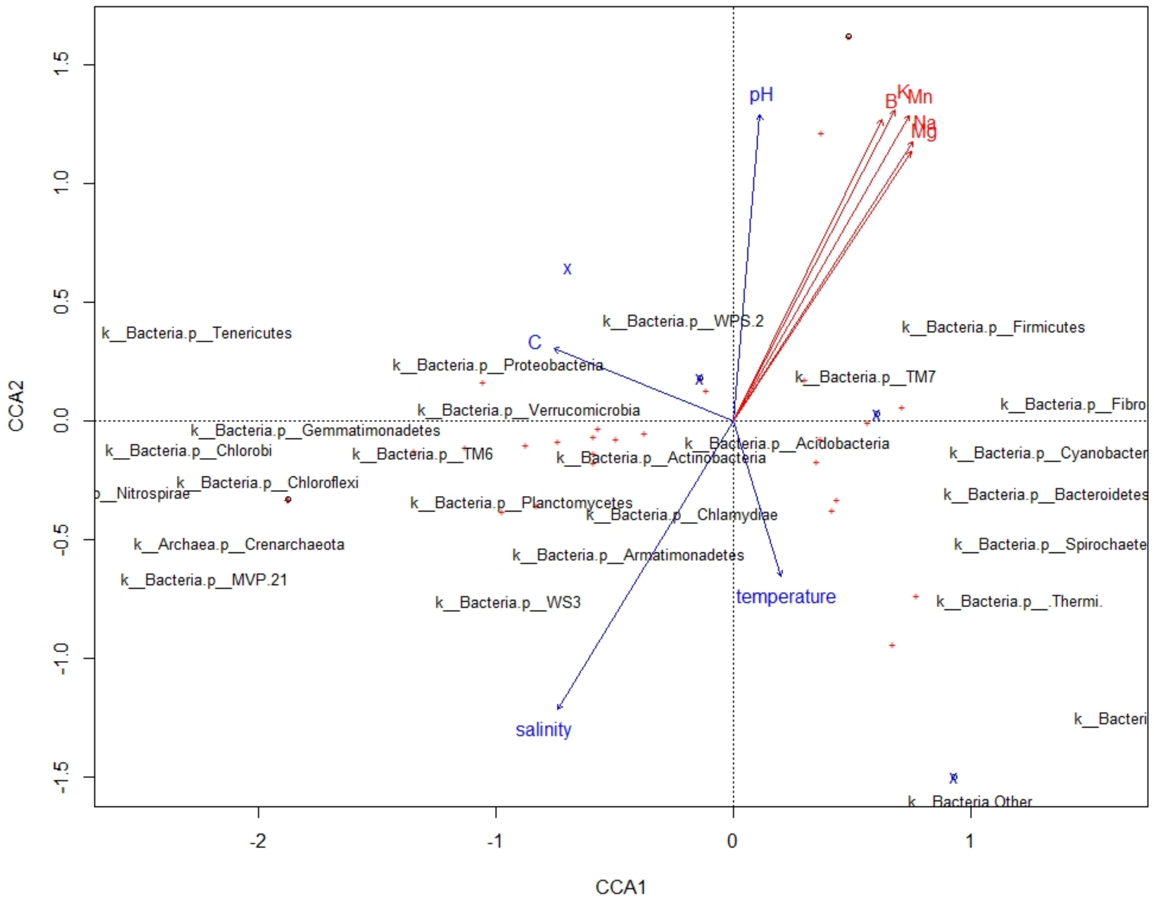


**Supplementary Figure 4:** CCA plot showing the relation of significant environmental factors with bacterial phylum. The blue arrows show statistically significant environmental factors. Red arrows show the remaining statistically insignificant environmental factors.

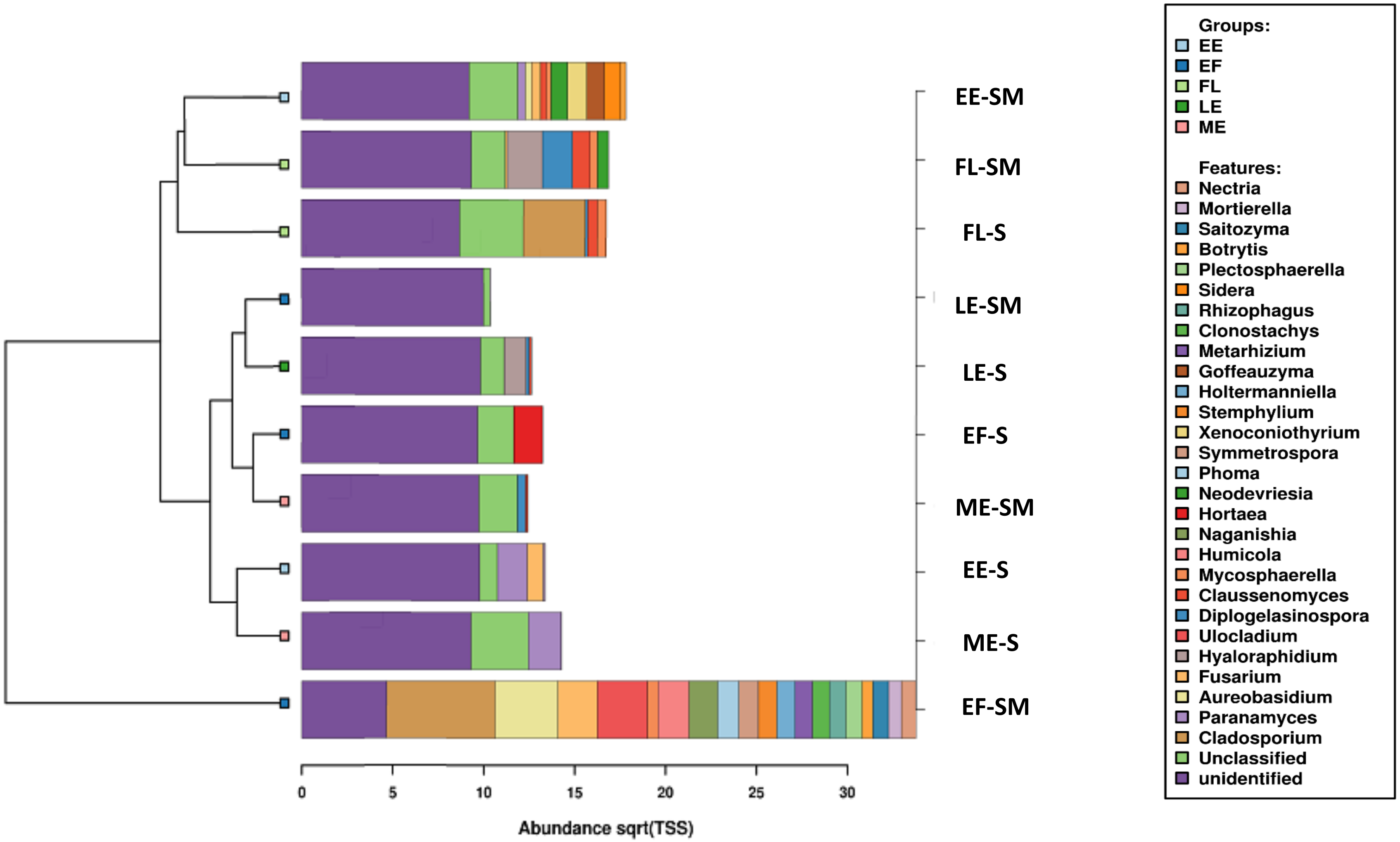


**Supplementary Figure 5:** Bar chart showing fungal composition at genera level for each layer at different time points. The samples have been arranged so more closely related are placed together.

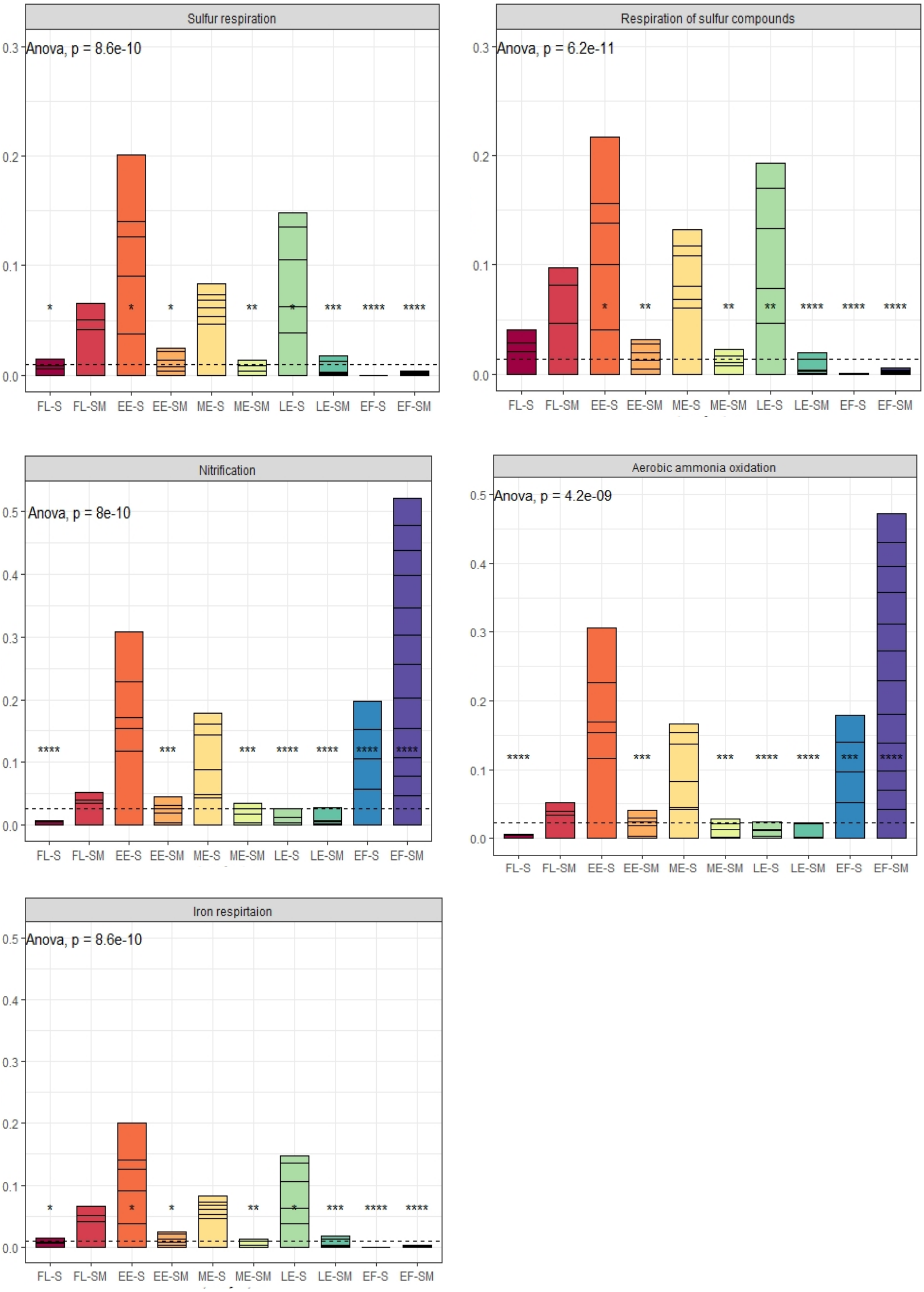


**Supplementary Figure 6:** Bar plots of individual functions of bacteria in the sediment and salt mat layer at different stages of Lake Magic. The statistical significance (ANOVA) is shown as asterisks.
